# Roles of sarcoplasmic reticulum Ca^2+^ ATPase pump in the impairments of lymphatic contractile activity in a metabolic syndrome rat model

**DOI:** 10.1101/2020.01.14.906503

**Authors:** Yang Lee, Sanjukta Chakraborty, Mariappan Muthuchamy

## Abstract

The intrinsic lymphatic contractile activity is necessary for proper lymph transport. Mesenteric lymphatic vessels from high-fructose diet-induced metabolic syndrome (MetSyn) rats exhibited impairments in its intrinsic phasic contractile activity; however, the molecular mechanisms responsible for the weaker lymphatic pumping activity in MetSyn conditions are unknown. Several metabolic disease models have shown that dysregulation of sarcoplasmic reticulum Ca^2+^ ATPase (SERCA) pump is one of the key determinants of the phenotypes seen in various muscle tissues. Hence, we hypothesized that a decrease in SERCA pump expression and/or activity in lymphatic muscle influences the diminished lymphatic vessel contractions in MetSyn animals. Results demonstrated that SERCA inhibitor, thapsigargin, significantly reduced lymphatic phasic contractile frequency and amplitude in control vessels, whereas, the reduced MetSyn lymphatic contractile activity was not further diminished by thapsigargin. While SERCA2a expression was significantly decreased in MetSyn lymphatic vessels, myosin light chain 20, MLC_20_ phosphorylation was increased in these vessels. Additionally, insulin resistant lymphatic muscle cells exhibited elevated intracellular calcium and decreased SERCA2a expression and activity. The SERCA activator, CDN 1163 increased phasic contractile frequency in the vessels from MetSyn, thereby, partially restored lymph flow. Thus, our data provide the first evidence that SERCA2a modulates the lymphatic pumping activity by regulating phasic contractile amplitude and frequency, but not the lymphatic tone. Diminished lymphatic contractile activity in the vessels from the MetSyn animal is associated with the decreased SERCA2a expression and impaired SERCA2 activity in lymphatic muscle.

## Introduction

Insulin resistance is one of the major causes of metabolic syndrome (MetSyn) or related metabolic disorders that are associated with an enormous health burden worldwide ^1^. MetSyn is now one of the most prevalent diseases globally and increases the risk for all causes of mortalities, including cardiovascular diseases ^1,2^. Clinical studies have established the link between obesity and lymphatic dysfunction, which is associated with increased susceptibility for developing lymphedema ^3–6^. Mice heterozygous for Prox1, a master lymphatic endothelial transcription factor, consistently develop adult onset obesity coupled with increased chyle accumulation in the thoracic cavity ^7,8^. In addition, these mice exhibited higher leptin and insulin levels ^8^, which are pathological determinant factors of insulin resistance suggesting a direct role of the lymphatic system in metabolic dysfunction. We have previously reported that a high-fructose-fed rat model of MetSyn presented a significant reduction in lymphatic pumping as a consequence of decreased phasic contractile frequency and impaired intrinsic lymphatic muscle force production ^9,10^. These findings have been corroborated in the obese mouse models that diminished pressure-induced frequency in collecting lymphatic vessels ^11^. We have also demonstrated that insulin resistance directly impaired cellular bioenergetics and decreased the relative levels of the regulatory molecule, myosin light chain 20 (MLC_20_) in lymphatic muscle cells (LMCs). However, the direct mechanisms that reduce lymphatic pumping activity in the MetSyn animals have not been completely understood.

The active spontaneous pumping of lymphatics is achieved by the intrinsic contractile activity of the lymphatic muscle cells in the wall of collecting lymphatic vessels that produces the rhythmic phasic contractions. The lymphatic muscle cells exhibit unique characteristics similar to both vascular smooth muscle and cardiac muscle cells ^12^. Like vascular smooth muscle, lymphatics show the contractile activity that is regulated by various vasoactive (e.g., substance p, endothelin-1, histamine, acetylcholine, etc.) and mechanical factors (e.g., transmural pressure, flow, etc.). In addition, lymphatic muscle displays a rapid phasic contraction that is mainly achieved by the intrinsic pumping characteristics. While the resting membrane potential is mediated by Cl^−^ ^13^ and voltage gated K^+^ channel ^14^, lymphatic contractions are predominantly regulated by Ca^2+^ influx ^15,16^. The intracellular Ca^2+^ concentration determines the lymphatic vessel contraction and similar to most other smooth muscle types, Ca^2+^ binds to calmodulin to form an active Ca^2+^/calmodulin complex, which activates myosin light chain kinase, a key regulatory molecule that phosphorylate MLC_20_ ^17–20^. Since MLC_20_ phosphorylation is increased in the insulin resistant LMCs^21^, we propose that regulatory molecules of Ca^2+^ and/or Ca^2+^ homeostasis are impaired in the MetSyn lymphatics.

The endoplasmic reticulum (ER) is the main storage site of intracellular Ca^2+^ that maintains intracellular Ca^2+^ levels ~10,000-fold lower than extracellular and ER Ca^2+^ concentrations ^22,23^. Re-uptake of Ca^2+^ into the ER by sarcoplasmic reticulum Ca^2+^- ATPase (SERCA) is necessary for muscle relaxation and restores ER Ca^2+^ levels for subsequent systolic and diastolic cycles followed by transiently increased intracellular Ca^2+^ levels. Alterations in Ca^2+^ homeostasis have been shown to trigger lymphatic dysfunction. When L-type Ca^2+^ channels were disrupted, stretch-induced lymphatic contractile amplitude was diminished; whereas, T-type, ‘transient,’ Ca^2+^ channel inhibition reduced the stretch-induced phasic contractile frequency in the lymphatics ^24^. Lymph flow was reduced by disrupting ER Ca^2+^ in bovine lymphatic vessels ^25^. Additionally, SERCA2 activity and expression are diminished in vascular smooth muscle ^26,27^ and heart ^28,29^ in different animal models of obesity/diabetes, highlighting a potential pathological role for SERCA2 dysfunction and disturbed intracellular Ca^2+^ homeostasis in the development of metabolic abnormalities in insulin resistance and diabetes. However, the role of SERCA2 in lymphatic pumping activity and possible pathophysiological roles in MetSyn have not yet been examined.

Our previous data showed negative chronotropic effects at all transmural pressures that effectively reduced the intrinsic flow generating capacity of mesenteric lymphatic vessels in MetSyn rats ^9,10^. Additionally, insulin resistance increased MLC_20_ phosphorylation in LMCs ^21^ that is mediated by intracellular Ca^2+^ ^20^. Hence, we hypothesized that a decrease in SERCA expression and/or activity in lymphatic muscle influences Ca^2+^ homeostasis in LMCs, and consequently, diminishes lymphatic contractile activity in MetSyn animals. To test this hypothesis, we assessed the expression of SERCA2 isoforms in lymphatic muscle and determined the role of SERCA2 in the regulation of lymphatic contraction in the normal and MetSyn conditions.

## Results

### MetSyn rats exhibit an altered body composition by decreasing skeletal muscle mass, but increasing body fat deposition and cardiac muscle mass

We have previously reported hyperinsulinemia and hyperlipidemia in high-fructose-induced MetSyn animals ^9^. As we expected, blood glucose levels in the high-fructose diet fed rats were significantly increased (7.37 ± 0.39 vs.13.37 ± 0.55mM, control vs. MetSyn, p<0.05) compared with the control group rats in the normal-diet chow (Table 1). As we reported in our previous studies, we did not observe any significant increase in body mass over the 7-10 week diet period in MetSyn rats comparing to the control group. However, skeletal muscle mass was found to be decreased in the MetSyn group when compared to the controls, which is consistent with the common pathological phenotype in metabolic disorders ^30^. Soleus and TA muscles from MetSyn rats showed a significant decrease in muscle mass (p<0.05) and in the fiber type distribution that is skewed to the smaller muscle fiber (Figures 1 A-D). In contrast, accumulation of body fat in both visceral and inguinal subcutaneous fat pad was significantly higher in MetSyn rats (Table 1). Additionally, heart weight was significantly increased in MetSyn animals and exhibited enlarged cardiac myocytes comparing with the myocytes from control rats (Table 1 and Figure 1).

**Figure 1.**
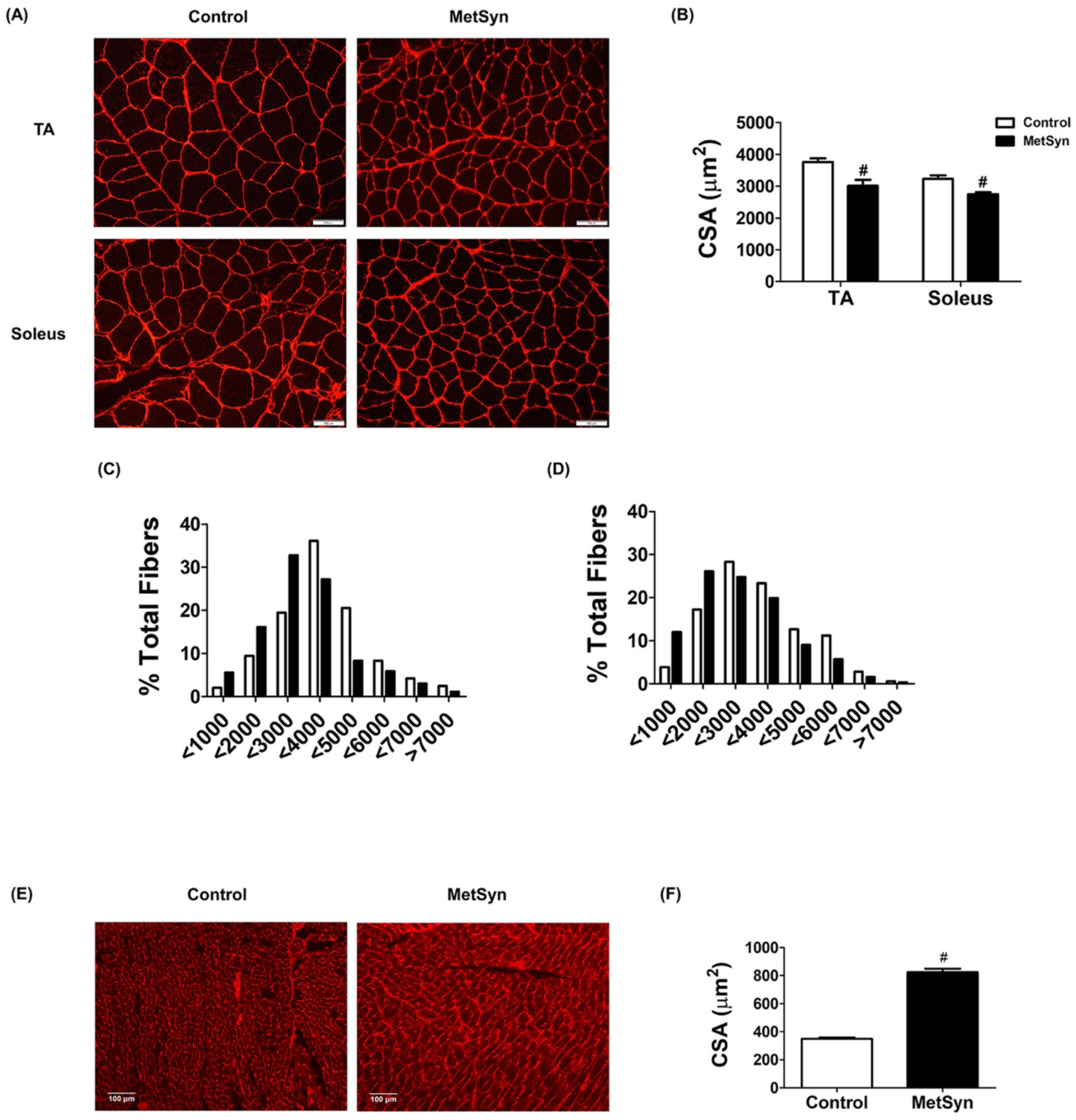
Skeletal muscle atrophy and cardiac hypertrophy in MetSyn animals. A) WGA staining shows skeletal muscle sarcolemma staining in TA (upper panel) and Soleus (lower panel) for control (left) and MetSyn (right). Image were obtained with x20 objective (NA = 0.7) on a fluorescence microscope. B) CSA of TA and soleus was quantified. Each individual myocyte was quantified and average cross sectional area was plotted (n=180 myocytes from 9 field of views from 3 animals/group). # indicates p<0.001 vs. control. C) TA CSA was displayed by fiber type distribution. D) Soleus CSA was displayed by fiber type distribution. E) WGA staining shows cardiac myocyte membrane staining. Images were obtained with x10 objective (NA=0.5). F) MetSyn increased cardiac myocyte CSA and quantified data was plotted (n=450 myocytes from 9 field of views from 3 animals/group). # indicates p<0.001 vs. control. Data are presented as mean ± SE.

**Table 1.** Effect of high fructose dietary feeding on MetSyn

| **Table 1. Effect of high fructose dietary feeding on MetSyn** |  |  |
| --- | --- | --- |
|  | Control (n=18) | MetSyn (n=18) |
| Body weight (g) | 390.42 ± 11.43 | 384.39 ± 14.83 |
| Glucose (mM) | 7.37 ± 0.39 | 13.37 ± 0.55 * |
| Visceral epididymal adipose fat (%) | 0.75 ± 0.05 | 1.74 ± 0.14* |
| Subcutaneous inguinal adipose fat (%) | 0.62 ± 0.06 | 1.47 ± 0.112* |
| Heart weight (g) | 1.29 ± 0.06 | 1.34.6 ± 0.04* |
| Heart/Bwt (mg/g) | 3.109 ± 0.09 | 3.48 ± 0.07* |
| Soleus (mg) | 218.1 ± 8.83 | 170.9 ± 8.54* |
| Soleus/Bwt (mg/g) | 0.56 ± 0.02 | 0.45 ± 0.038* |
| TA (mg) | 849.9 ± 44.58 | 684 .2 ± 57.86* |
| TA/Bwt (mg/g) | 2.19 ± 0.07 | 1.78 ± 0.15* |

Values are expressed as means ± SE. \*p<0.05

### SERCA inhibition abolishes intrinsic lymphatic pumping activity

To assess the role of SERCA in the regulation of lymphatic vessel contractility, we examined lymphatic vessel contractions in different transmural pressures in the absence or presence of various doses of SERCA pump inhibitor, thapsigargin. At concentrations higher than 5μM, thapsigargin completely inhibited lymphatic vessels phasic contractions (data not shown). Therefore, we examined the effects of thapsigargin in the dose range of 500nM to 2μM on lymphatic vessel contractile properties. While 500nM dose of thapsigargin did not affect the contractile frequency, amplitude, ejection fraction and fractional pump flow at any tested transmural pressures, 1.0μM thapsigargin significantly reduced the contraction amplitude and the ejection fraction at P=3 or P=5 cmH_2_O (Figures 2 B-E). Though the contractile frequency of the lymphatics was not significantly reduced when treated with 1.5μM of thapsigargin, the pressure-dependent increases in contractile frequencies were blunted (11.73 ± 1.71, 12.3 ± 1.7, 1.37 ± 2.5 vs. 8.09 ± 1.61, 8.26 ±1.45, 8.24 ± 1.33 contractions/min at p=1, 2, and, 3 cmH_2_O, control vs. 1.5μM thapsigargin respectively). Additionally, it significantly reduced ejection fraction, contractile amplitude, and fractional pump flow (p<0.05, Figure 2). 2μM thapsigargin was found to severely impair the lymphatic contractions with decreased frequency and amplitude, and thus fractional pump flow and ejection fraction (Figure 2). Therefore, we employed 1.5μM of thapsigargin to examine the role of SERCA activity in the regulation of lymphatic vessel contractility of MetSyn animals.

**Figure 2.**
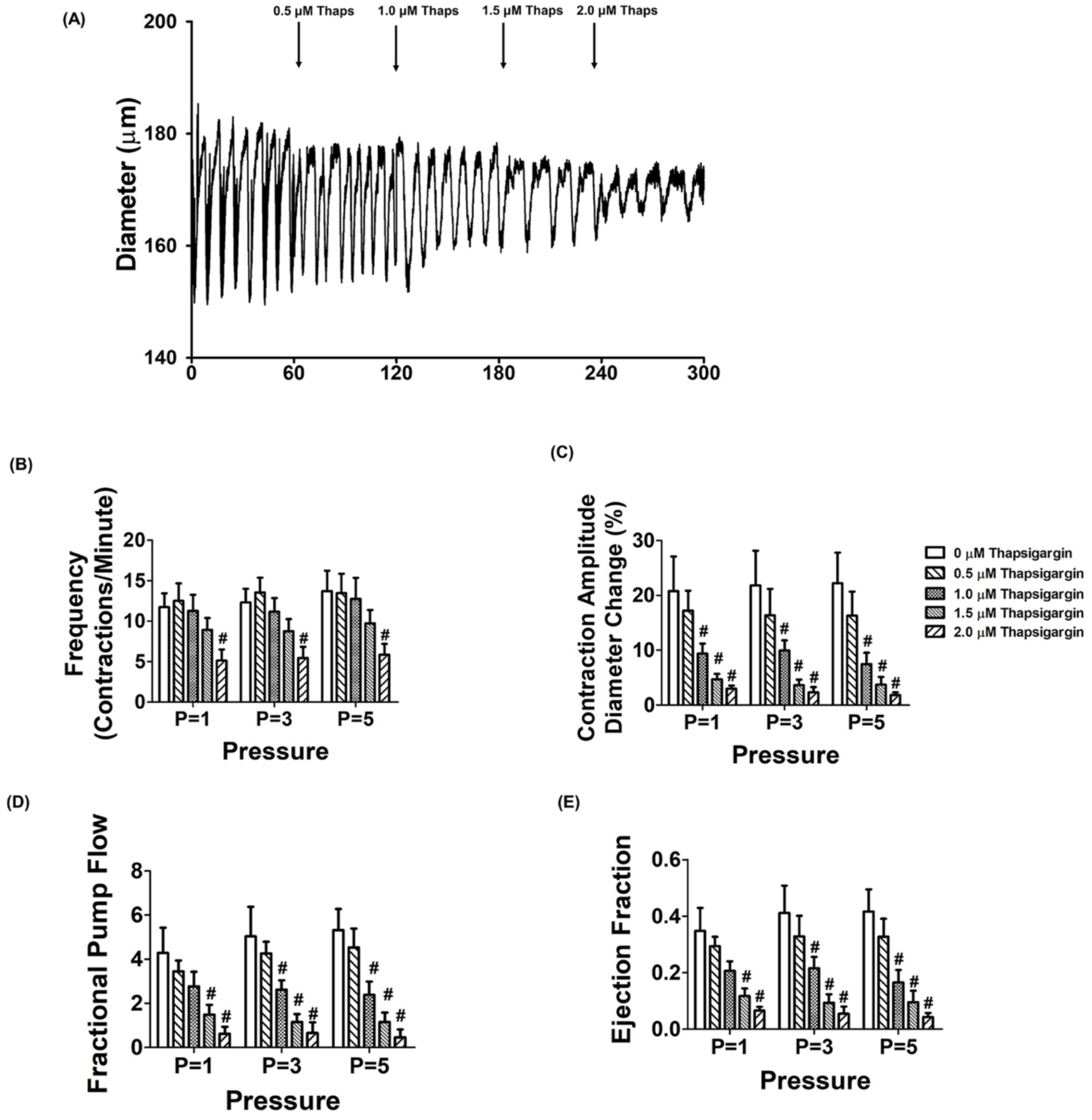
Inhibition of SERCA with thapsigargin causes impaired lymphatic pumping of rat mesenteric collecting lymphatic vessels. A) An example of the lymphatic diameter over time before and after treatment with increasing concentrations of the SERCA inhibitor thapsigargin at 3cmH_2_O. The SERCA inhibitor, thapsigargin significantly decreased B) contractile frequency, C) contractile amplitude, D) fractional pump flow, and E) ejection fraction (n=6 lymphatic vessels from 5 animals). #<0.05 vs. control (no thapsigargin). Data are presented as mean ± SE.

### SERCA activity is diminished in MetSyn lymphatic vessels

In control lymphatic vessels, as discussed above, the phasic contractile frequency was decreased at each employed pressure, in the presence of thapsigargin (10.27 ± 0.59 vs. 7.96 ± 0.9, 11.12 ± 0.69 vs. 9.14 ± 1.27, and 12.19 ± 0.73 vs. 9.58 ± 1.06 contractions per min, at 1, 2, and 3 H_2_O pressure respectively), but was not statistically significant (Figures 3A and C, Table 2). In MetSyn animals, contractile frequency was significantly decreased compared to control animals (6.59 ± 0.8, 7.67 ± 1.02, 8.96 ± 0.89 at P=1, 2, and 3 cmH_2_O respectively, p<0.05, Figures 3B and C, Table 2), as we have previously reported ^9,10^. However, thapsigargin did not further decrease the contractile frequency of lymphatic vessels from MetSyn animals. In addition, the reduction in contractile frequency due to thapsigargin in control animals was found to be similar to the contractile frequency in MetSyn animals (Figure 3C). Additionally, thapsigargin significantly reduced the ejection fraction in both control and MetSyn animals at transmural pressures, P=1, 3 and 5 cmH_2_O, while there was no significant difference between the control and MetSyn groups (Figure 3D). The fractional pump flow and lymph flow were also reduced significantly by thapsigargin in control lymphatic vessels in all transmural pressures (p<0.05, Figures 3 E & F). While fractional pump flow and lymph flow were significantly decreased in the lymphatic vessels from MetSyn rats compared to the control group, thapsigargin did not further diminish these parameters in MetSyn group (Figures 3 E & F). A summary of all the lymphatic contractile parameters is given in Table 2.

**Figure 3.**
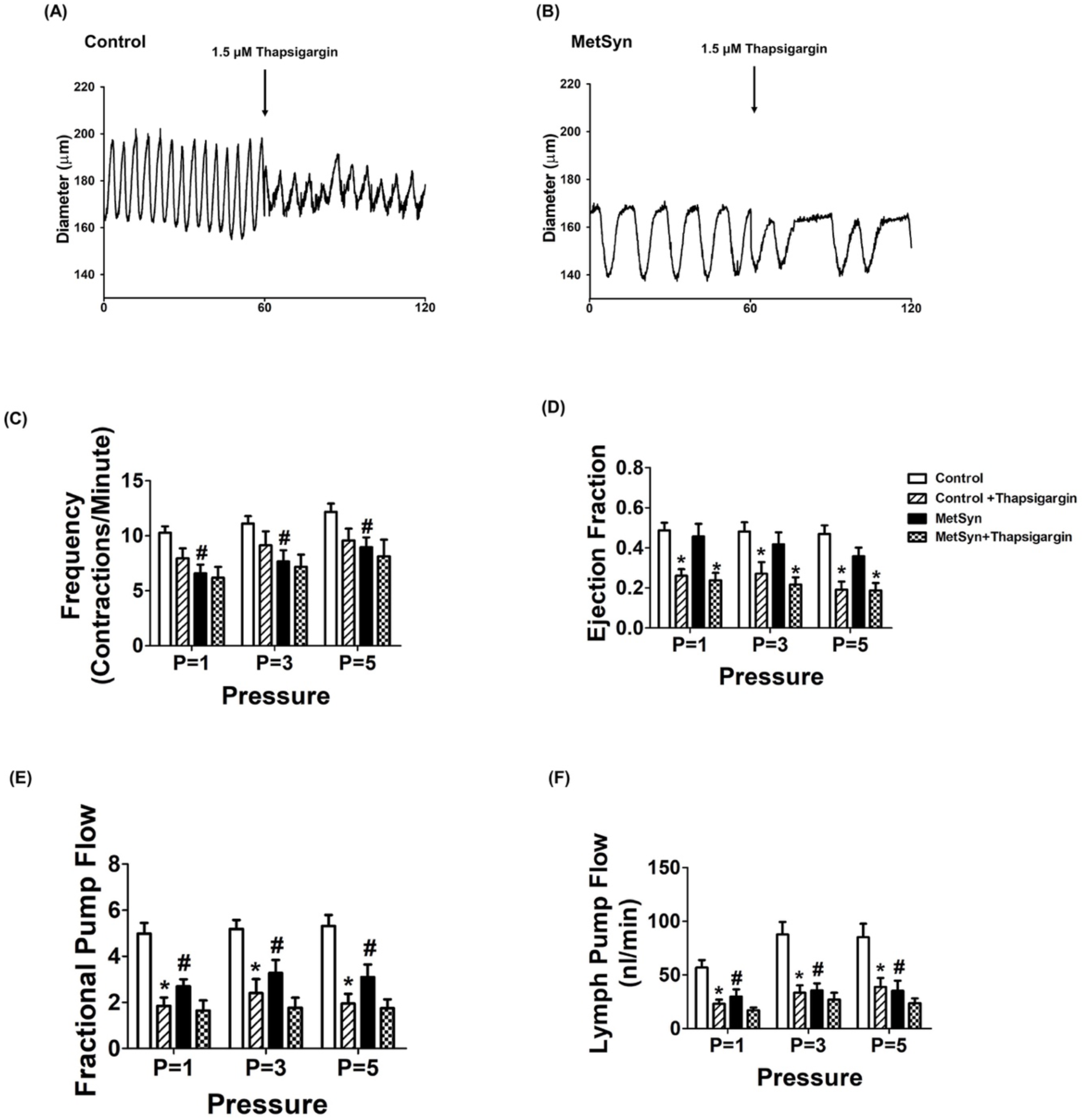
Lymphatic pump dysfunction in isolated lymphatic vessels from MetSyn animals was due to impaired SERCA activity. Representative diameter traces over 1 minute period of: A) control and B) MetSyn lymphatic vessel at pressure 3 cmH_2_O in the presence of SERCA inhibitor. The effect of SERCA inhibitor on: C) contractile frequency, D) ejection fraction, E) fractional pump flow, and F) lymph flow, for control (n=12 from 6 animal) and MetSyn (n=10 from 6 animal). # indicates p<0.05 vs. control groups at any value of transmural pressure. * indicates p<0.05 vs. each cohort controls at any value of transmural pressure. Data are presented as mean ± SE.

**Table 2.** Contractile parameters for control and MetSyn mesentery lymphatic vessels presence and absence of thapsigargin (1.5μm) or CND1163 (5 μM)

| Parameters | Cohort | P=1 | P=3 | P=5 |
| --- | --- | --- | --- | --- |
| Outer diastolic diameter (µm) | Control | 136.53 ± 6.38 | 152.14 ± 7.1 | 161.58 ± 7.21 |
|  | Control Thapsigargin | 133.71 ± 4.28 | 151.22 ± 7.01 | 160.5 ± 7.53 |
|  | MetSyn | 127.8 ± 6.43 | 135.02 ± 6.92 | 141.95 ± 7.75 |
|  | MetSyn Thapsigargin | 121.37 ± 6.12 | 132.89 ± 6.86 | 141.39 ± 6.89 |
| Contraction frequency per minute | Control | 10.27 ± 0.59 | 11.12 ± 0.69 | 12.19 ± 0.73 |
|  | Control Thapsigargin | 7.96 ± 0.9 | 9.14 ± 1.27 | 9.58 ± 1.06 |
|  | MetSyn | **6.59 ± 0.8<sup>#</sup>** | **7.67 ± 1.02<sup>#</sup>** | **8.96 ± 0.89<sup>#</sup>** |
|  | MetSyn Thapsigargin | 6.17 ± 1.09 | 7.16 ± 1.19 | 8.13 ± 1.44 |
| Ejection fraction | Control | 0.49 ± 0.04 | 0.48 ± 0.05 | 0.47 ± 0.04 |
|  | Control Thapsigargin | **0.26 ± 0.03\*** | **0.27 ± 0.06\*** | **0.19 ± 0.04\*** |
|  | MetSyn | 0.45 ± 0.06 | 0.41 ± 0.06 | 0.36 ± 0.04 |
|  | MetSyn Thapsigargin | **0.24 ± 0.04\*** | **0.22 ± 0.03\*** | **0.18 ± 0.03\*** |
| Lymph pump flow (nL/min) | Control | 57.0 ± 6.96 | 87.73 ± 11.64 | 85.17 ± 12.63 |
|  | Control Thapsigargin | **23.2 ± 3.71\*** | **33.45 ± 6.88\*** | **38.83 ± 8.41\*** |
|  | MetSyn | **29.82 ± 6.68<sup>#</sup>** | **35.52 ± 6.74<sup>#</sup>** | **35.24 ± 9.5<sup>#</sup>** |
|  | MetSyn Thapsigargin | 17.15 ± 2.51 | 26.96 ± 6.47 | 23.63 ± 4.51 |
| Fractional pump flow (min<sup>-1</sup>) | Control | 4.98 ± 0.47 | 5.12 ± 0.38 | 5.32 ± 0.47 |
|  | Control Thapsigargin | **1.85 ± 0.35\*** | **2.40 ± 0.61\*** | **1.95 ± 0.42\*** |
|  | MetSyn | **2.70 ± 0.30<sup>#</sup>** | **3.28 ± 0.56<sup>#</sup>** | **3.09 ± 0.54<sup>#</sup>** |
|  | MetSyn Thapsigargin | 1.64 ± 0.43 | 1.77 ± 0.43 | 1.75 ± 0.38 |
| Contraction amplitude (%) | Control | 23.51 ± 3.28 | 24.01 ± 2.2 | 22.05 ± 2.18 |
|  | Control Thapsigargin | **12.20 ± 1.79\*** | **14.40 ± 2.37\*** | **11.85 ± 2.13\*** |
|  | MetSyn | 18.56 ± 3.77 | 19.13 ± 3.58 | 19.27 ± 3.16 |
|  | MetSyn Thapsigargin | 10.29 ± 1.51 | 9.93 ± 1.67 | 11.37 ± 2.01 |
| Tonic index | Control | 14.14 ± 2.82 | 9.52 ± 2.36 | 7.62 ± 2.55 |
|  | Control Thapsigargin | 15.57 ± 2.42 | 10.3 ± 1.57 | 8.63 ± 2.01 |
|  | MetSyn | 14.11 ± 1.94 | 10.72 ± 1.94 | 9.64 ± 1.78 |
|  | MetSyn Thapsigargin | 18.03 ± 4.19 | 12.22 ± 3.62 | 10.96 ± 3.27 |
| Outer diastolic diameter (µm) | Control | 150.2 ± 17.93 | 164.7 ± 11.52 | 171.6 ± 51 |
|  | Control CDN1163 | 143.2 ± 17.05 | 162.8 ± 13.7 | 171.2 ± 11.06 |
|  | MetSyn | 137.8 ± 11.94 | 146.1 ± 11.44 | 151.1 ± 10.54 |
|  | MetSyn CDN1163 | 134.2 ± 10.44 | 142.0 ± 9.78 | 148.2 ± 8.92 |
| Contraction frequency per minute | Control | 11.83 ± 1.71 | 12.12 ± 1.48 | 12.29 ± 1.33 |
|  | Control CDN1163 | 15.82 ± 1.8 | **19.83 ± 2.94\*** | **21.68 ± 4.08\*** |
|  | MetSyn | **5.32 ± 0.71<sup>#</sup>** | **6.03 ± 0.59<sup>#</sup>** | **7.08 ± 0.52<sup>#</sup>** |
|  | MetSyn CDN1163 | **14.86 ± 3.41\*** | **16.68 ± 4.02\*** | **17.24 ± 3.87\*** |
| Ejection fraction | Control | 0.38 ± 0.37 | 0.42 ± 0.06 | 0.39 ± 0.05 |
|  | Control CDN1163 | 0.29 ± 0.39 | 0.31 ± 0.07 | 0.28 ± 0.07 |
|  | MetSyn | 0.30 ± 0.46 | 0.37 ± 0.06 | 0.32 ± 0.05 |
|  | MetSyn CDN1163 | 0.22 ± 0.51 | 0.20 ± 0.03 | 0.19 ± 0.03 |
| Lymph pump flow (nL/min) | Control | 86.31 ± 21.66 | 104.47 ± 16.84 | 105.32 ± 21.42 |
|  | Control CDN1163 | 68.39 ± 22.09 | 78.48 ± 12.94 | 70.72 ± 19.9 |
|  | MetSyn | 43.79 ± 13.42 | **46.5 ± 11.39<sup>#</sup>** | **50.92 ± 8.62<sup>#</sup>** |
|  | MetSyn CDN1163 | 36.86 ± 4.72 | 39.42 ± 5.35 | 38.62 ± 6.57 |
| Fractional pump flow (min<sup>-1</sup>) | Control | 4.49 ± 0.41 | 4.94 ± 0.72 | 4.62 ± 0.98 |
|  | Control CDN1163 | 5.29 ± 1.34 | 5.49 ± 1.25 | 5.27 ± 1.66 |
|  | MetSyn | **2.13 ± 0.73<sup>#</sup>** | **2.16 ± 0.68<sup>#</sup>** | **2.19 ± 0.57<sup>#</sup>** |
|  | MetSyn CDN1163 | 3.31 ± 0.69 | 3.14 ± 0.60 | 2.82 ± 0.64 |
| Contraction amplitude (%) | Control | 25.50 ± 4.03 | 29.05 ± 3.15 | 26.13 ± 2.82 |
|  | Control CDN1163 | 16.66 ± 3.65 | 15.8 ± 5.4 | 24.72 ± 6.47 |
|  | MetSyn | 23.37 ± 5.9 | 21.66 ± 3.85 | 21.00 ± 4.19 |
|  | MetSyn CDN1163 | 16.33 ± 2.34 | 14.77 ± 2.28 | 13.71 ± 2.29 |
| Tonic index | Control | 15.38 ± 1.01 | 9.25 ± 0.79 | 7.19 ± 0.99 |
|  | Control CDN1163 | 19.04 ± 2.38 | 10.43 ± 2.03 | 7.29 ± 1.75 |
|  | MetSyn | 14.29 ± 0.51 | 9.62 ± 1.27 | 7.83 ± 0.86 |
|  | MetSyn CDN1163 | 16.06 ± 2.36 | 11.76 ± 1.86 | 9.27 ± 1.56 |

Values are expressed as mean ± SE. \* indicate significant difference between Thapsigargin or CDN1163 treatment within each cohort group at any value of transmural pressure (p<0.05). # indicate significant difference between control and MetSyn lymphatic vessels in control group at any value of transmural pressure (p<0.05).

### Decreased SERCA2a expression in MetSyn lymphatic vessel

To assess the expression of SERCA, we performed immunofluorescence analyses on isolated mesenteric lymphatic vessels from control and MetSyn rats using striated muscle-specific SERCA2a or striated and smooth muscle isoform, SERCA2b specific antibody co-stained with α-smooth muscle actin (α-SMA) antibody. Both SERCA2a and SERCA2b positive staining were revealed on the lymphatic vessel wall. SERCA2a and SERCA2b are co-stained with α-SMA in lymphatic muscle, indicating both SERCA2a and SERCA2b are present in lymphatic muscle (Figures 4A and C). Negative controls were performed with normal rabbit IgG (data not shown). Further quantitative analyses showed that SERCA2a expression was significantly decreased in the MetSyn lymphatic muscle compared to control group (0.56 ± 0.39 fold, p<0.001, n=9 vessels from three animals/group, Figures 4A and B); however, there was no significant differences in the SERCA2b expression among the control and MetSyn groups. We further examined the relative levels of MLC_20_ phosphorylation in the control and MetSyn lymphatic vessels. MetSyn lymphatic vessels displayed significantly higher levels of phosphorylated MLC_20_ (p<0.008, Figure 4E-F).

**Figure 4.**
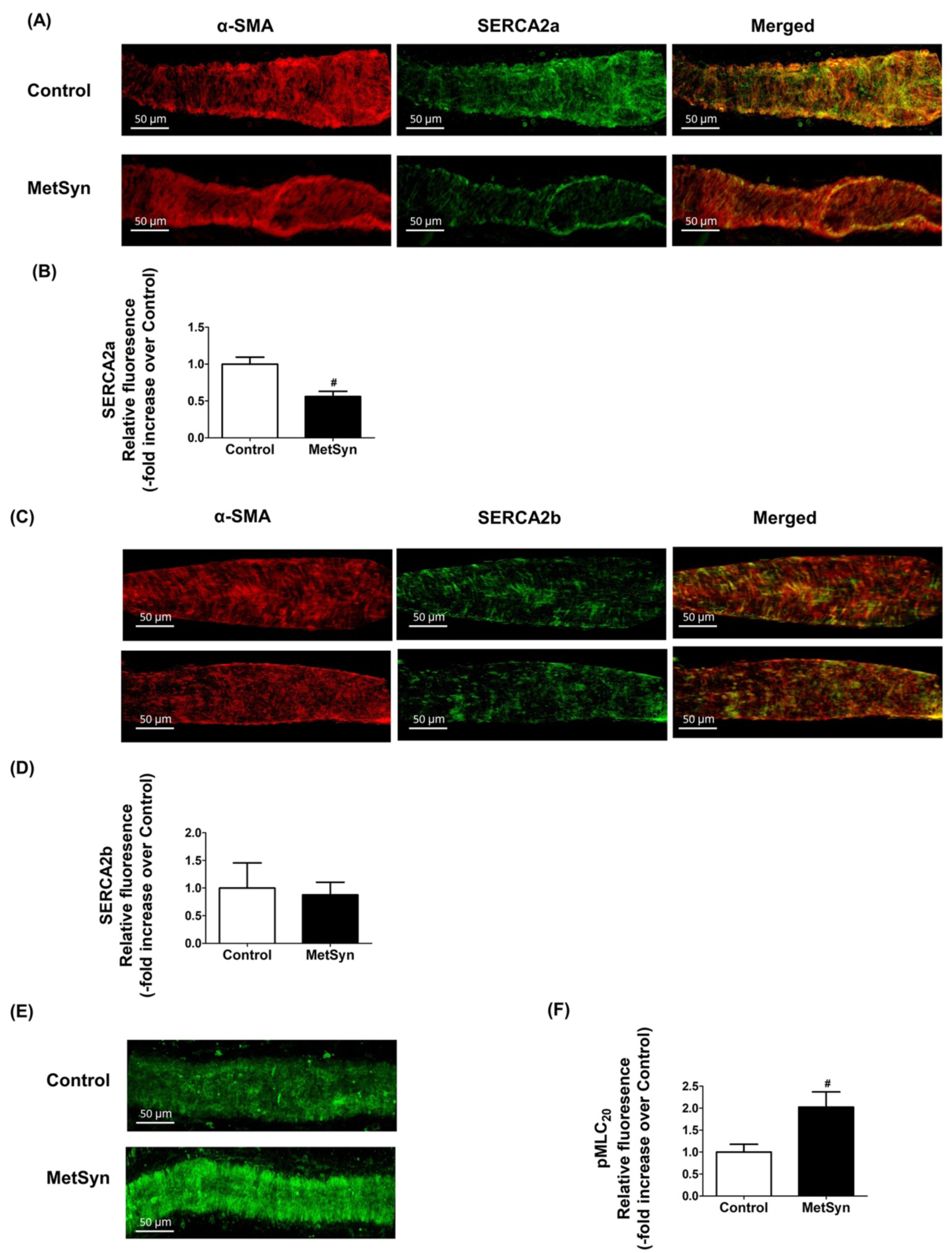
Decreased SERCA2a expression coupled with elevation of MLC_20_ phosphorylation in lymphatic vessels from MetSyn animals. A) Representative images of SERCA2a and α-SMA staining for lymphatic vessels from control and MetSyn rats. Images were obtained with x 40 objective (NA =0.9) on a confocal microscope. Scale bars, 50μm. B) Average projections were quantified and plotted for control (n = 18 fields of views from 6 vessels from 3 animals) and MetSyn (n=30 fields of views from 6 vessels from 3 animals). Quantification of SERCA2a was plotted. # indicates p<0.001 vs control. C) Representative images of SERCA2b and α-SMA staining for lymphatic vessels from control and MetSyn rats. Images were obtained with x40 objective (NA=0.9) on a confocal microscope, and average projections were presented. Scale bars, 50μm. D) Images were quantified and plotted for control (n = 9 fields of views from 4 vessels from 2 animals) and (n= 10 fields of view from 4 vessels from 2 animals). E). Representative images of MLC_20_ phosphorylation staining for lymphatic vessels from control and MetSyn rats. Images were obtained with x40 objective (NA=0.9) on a confocal microscope. Scale bars, 50μm. F). Average projections were quantified and plotted for control (n=17 fields of view from 6 vessels from 3 animals) and MetSyn (n=12 fields of view from 6 vessels from 3 animals). # p<0.008 vs. Control. Data are presented as mean ± SE.

### Insulin resistance impaired SERCA2 activity in LMCs

To address whether insulin resistance conditions in LMCs impair SERCA activity and expression, and calcium regulation, LMCs were cultured in hyperglycemia and hyperinsulinemia conditions for 48 h as described in our previous study ^21^. Insulin resistant LMCs showed elevated basal intracellular Ca^2+^ levels (92.1 ± 2.64nM) compared to controls (77.19 ± 1.01nM, p<0.001) and other groups (Figures 5 A and B). We used 5μM of thapsigargin that prevented Ca^2+^ uptake into the endoplasmic reticulum by blocking SERCA. Thus, inhibiting SERCA contributed to significantly increase peak intracellular Ca^2+^ in LMCs. Peak intracellular Ca^2+^ and transient time to reach the peak Ca^2+^ levels were not significantly different between the groups (Figure 5A). The amplitude between basal and peak Ca^2+^ levels in the presence of the thapsigargin was found to be significantly decreased in insulin resistant LMCs (Control, 52.36 ± 2.59nM vs. HG + Insulin, 30.52 ± 2.42nM, p<0.001), indicating decreased SERCA activity in these cells (Figures 5A and C). No difference was observed between groups in the peak Ca^2+^ levels when we depolarized the cells using 80mM K^+^ (data not shown). ER specific SEC61 protein staining showed no differences in the ER morphology between the groups. SERCA2a expression was significantly lower in insulin resistant LMCs when compared to control LMCs (−0.41 fold vs. control, p<0.032) (Figures 5D and E). Corroborating our immunofluorescence data, western blot analyses demonstrated that SERCA2a protein expression was decreased in insulin resistant LMCs (−0.58 fold vs. control, p<0.026) while SERCA2b expression remained unchanged (Figures 5F and G). Next, we tested whether increased extracellular free calcium level in LMCs would directly influence MLC_20_ phosphorylation levels as external calcium levels affect intracellular calcium level and muscle contractility ^31–33^. LMCs were grown in the media containing different free calcium concentration solution (calcium free to pCa3.5). MLC_20_ phosphorylation was found to be increased in LMCs grown under increasing extracellular Ca^2+^ concentrations and showed significant increase at pCa 6.5 or at higher pCa (p<0.05, Figure 5H).

**Figure 5.**
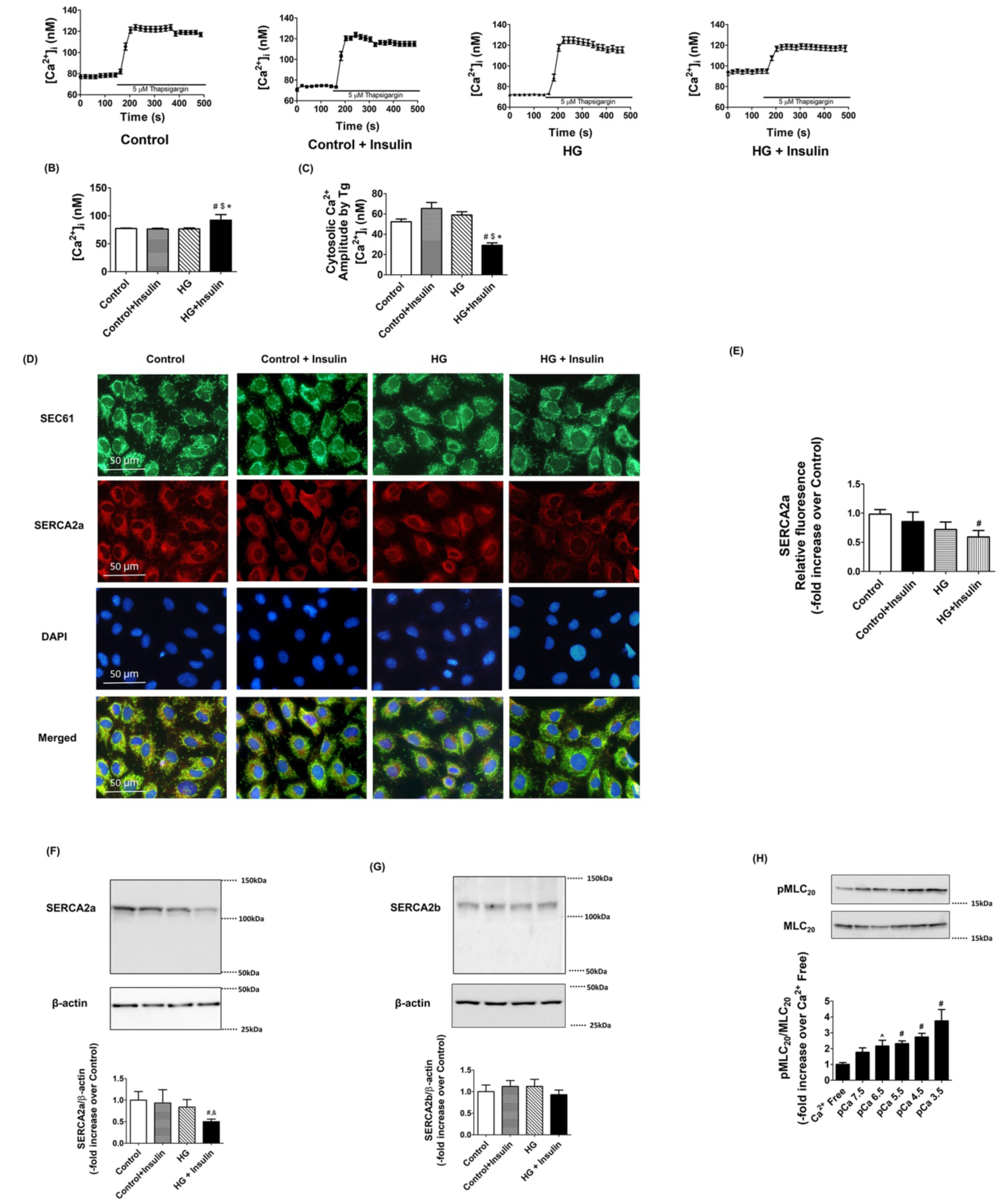
Insulin resistant conditions impaired SERCA activity and decreased SERCA2a expression in LMCs. A) An example tracing shows that the SERCA inhibitor, thapsigargin increased intracellular Ca^2+^ levels in the absence or presence of high glucose and insulin (n=100 cells from 5 different cultured LMC dishes/groups). B) Basal calcium level, before thapsigargin, was quantified and plotted. # indicates p<0.001 vs. Control. $ indicates p<0.001 vs. Control + Insulin. * indicates p<0.001 vs. HG. C) Amplitude between peak intracellular Ca^2+^ after thapsigargin and basal was quantified for determining SERCA activity and plotted. # indicates p<0.001 vs. Control. $ indicates p<0.001 vs. Control + Insulin. * indicates p<0.001 vs. HG. D). Representative immunofluorescence images of lymphatic muscle cells stained with antibodies specific for the ER (SEC61, green), SERCA2a (Red), and DAPI (blue). Images were obtained using x40 objective (NA=1.3) on a fluorescence microscope (n=9 field of views from 3 cultured dishes/group). Scale bars, 50μm. E). SERCA2a relative fluorescence intensity was quantified and plotted. # indicates p<0.032 vs. Control. Representative Western blots of SERCA2a (F) and SERCA2b (G) relative levels in LMCs treated with HG, insulin, or both together for 48h. The relative expression of SERCA2a/β-actin and SERCA2b/β-actin were quantified and plotted (n=3/group). # indicates p<0.05 vs. Control. $ indicates p<0.05 vs. Control + Insulin. H). Representative Western blots show increased free extracellular calcium concentration induced MLC_20_ phosphorylation in LMCs. The relative expression of pMLC_20_/total MLC_20_ (n=4/group). ^ indicates p<0.1 compared to Ca^2+^ free control. # indicates p<0.05 vs. Ca^2+^ free control. Data are presented as mean ± SE.

### Activation of SERCA pump partially improves lymphatic contractile activity in MetSyn animals

CDN1163 is a small molecule that activates SERCA2 by directly binding to the SERCA2 structure and increases SERCA2 V_max_ activity ^34–36^. In this study, we tested whether exogenously adding CDN1163 in the isolated lymphatic vessel preparations of MetSyn rats would improve its pumping activity. We selected different doses of CDN1163 (1, 5, and 10μM) based on previous studies. We initially determined the effects of CDN1163 on lymphatic contractile parameters of the control vessels. While lymphatic contractile frequency was not affected in the presence of 1μM CDN 1163, 5μM CDN1163 significantly increased contractile frequency of lymphatic vessels at transmural pressures, 3cmH_2_O (17.74 ± 2.7 contractions/min, p<0.022) and 5cmH_2_O (18.16 ± 1.9 contractions/min, p<0.016, Figure 6B). Lymphatic contractile frequency was increased at all different transmural pressures in the presence of 10μm CDN1163 (p<0.001, Figure 6B). Though CDN1163 did not alter significantly the contractile amplitude, stroke volume, lymph flow, tone, and ejection fraction, there was a trend that lymphatic contractile amplitude and ejection fraction were decreased in a dose dependent manner (data not shown). Therefore, even if CDN1163 significantly increased contractile frequency of control lymphatic vessels, lymphatic fractional pump flow did not change at all different transmural pressures in the presence of 5μm CDN1163 (Figure 6C). Hence, we used 5μm CDN1163 that showed positive chronotropic effect without negative inotropic effect (Figure 6A-C), to test whether CDN1163 ameliorates MetSyn induced impaired lymphatic contractile activity. CDN1163 significantly improved contractile frequency of lymphatic vessels from MetSyn animals (Figures 6 D, E and F). Similar to lymphatic vessels from control animals, SERCA activation did not improve fractional pump flow significantly in the lymphatic vessels from MetSyn group (Figure 6G). However, the decreased fractional pump flow in MetSyn vessels was not reduced significantly after treatment with CDN1163, indicating the partial protective effect of SERCA activator. The lymphatic contractile parameters of control and MetSyn rats are detailed in Table 2.

**Figure 6.**
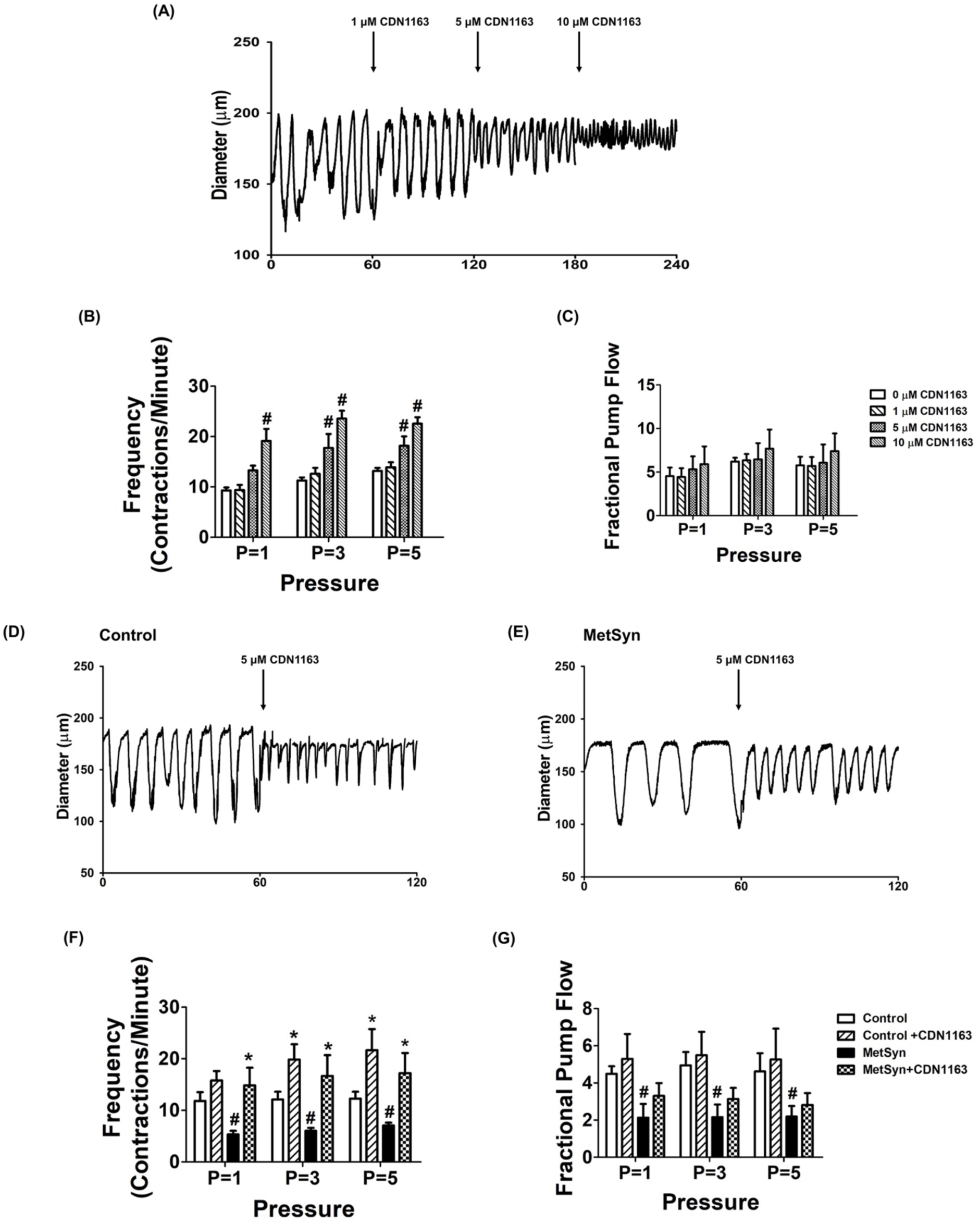
The effects of SERCA activator, CDN1163 on contraction frequency and lymph flow in isolated lymphatic preparations from control and MetSyn rats. A). Representative trace of control mesenteric lymphatic vessel at 3 cmH_2_O transluminal pressure with different doses of CDN1163 (n=5 lymphatic vessels). The effects of CND1163 on: B) contraction frequency and C) fractional pump flow. # indicates p<0.01 vs. control. Representative diameter traces of control (D) and MetSyn (E) mesenteric lymphatic vessels at pressure 3 cmH_2_O in the presence of CDN1163. The effects of CDN1163 on: F) contractile frequency and G) fractional pump flow (n=6 from 4 animal/group). # indicates p<0.05 vs. control groups at any value of transmural pressure. * indicates p<0.05 compared to each cohort controls at any value of transmural pressure. Data are presented as mean ± SE.

## Discussion

The data presented in this study provide the first evidence that the striated muscle-specific, SERCA2a pump that is present in lymphatic muscle modulates the lymphatic pumping activity by regulating phasic contractile amplitude and frequency, but not the lymphatic tone. Additionally, diminished lymphatic contractile activity in the vessels from the MetSyn animal is associated with the decreased SERCA2a expression and impaired SERCA2 activity. SERCA activator, CDN1163 significantly improved the contractile frequency and partially restored fractional pump flow in MetSyn mesenteric lymphatic vessels. Additionally, our data demonstrate that reduced SERCA2a expression resulted in impaired Ca^2+^ homeostasis in insulin resistant LMCs. Collectively, these data suggest MetSyn conditions diminished SERCA2a expression and SERCA activity in lymphatic muscle, consequently reduced lymphatic contractile activity in MetSyn rats (Figure 7).

**Figure 7.**
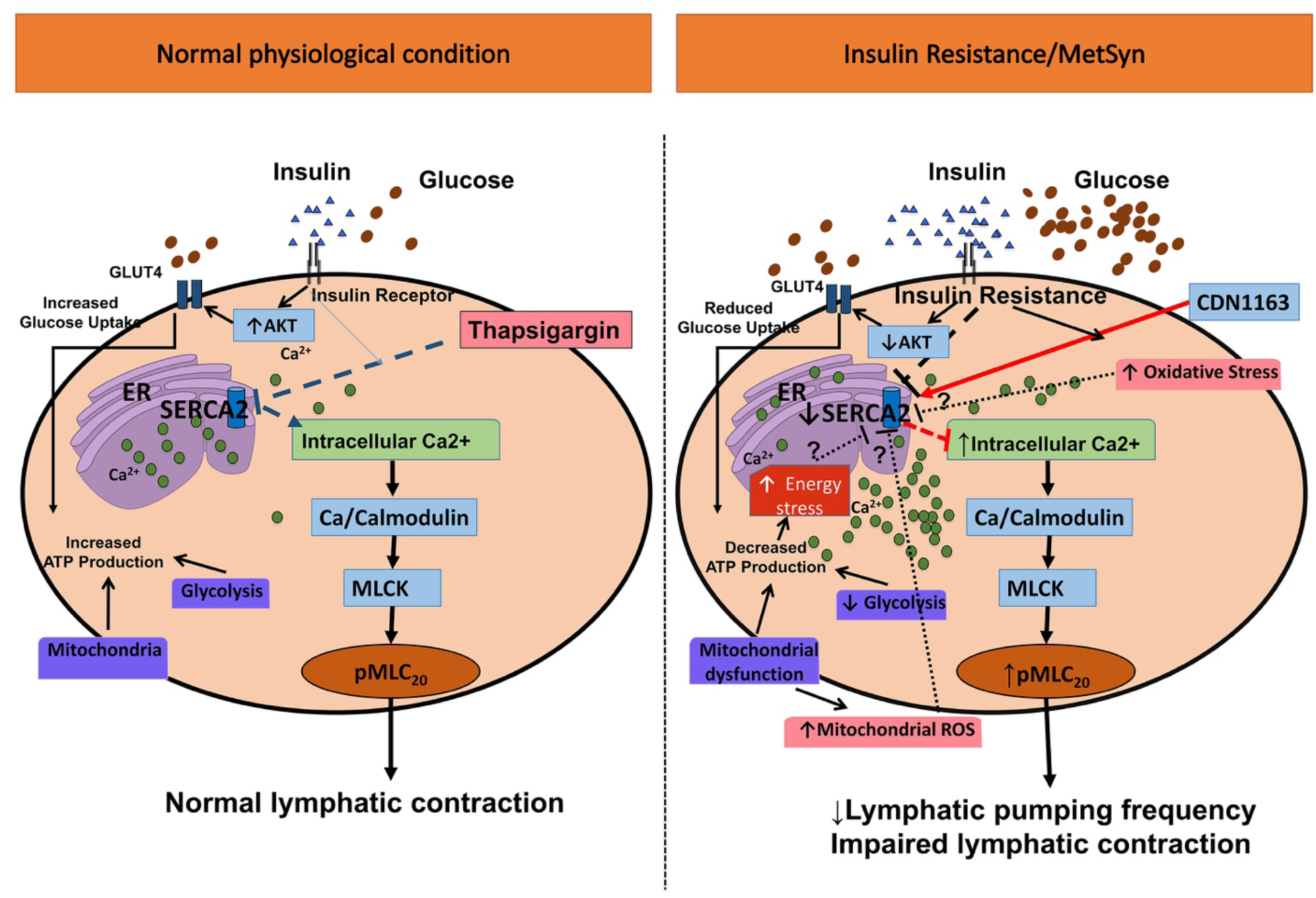
Schematic representation of lymphatic vessel contraction under normal and MetSyn/insulin resistant conditions. In normal physiological condition, Ca^2+^ homeostasis is critical for lymphatic pump regulation and proper lymph flow. Inhibition of SERCA directly diminishes lymphatic contraction and lymph flow. Insulin resistance or high fructose diet-induced MetSyn condition impairs SERCA2a expression and activity, which disturbs intracellular Ca^2+^ homeostasis. Elevated intracellular Ca^2+^ promotes MLC_20_ phosphorylation via Ca^2+^/calmodulin and MLCK that results in impaired lymphatic contractile activity and lymph flow in MetSyn animals.

In keeping with previous studies, we found that the MetSyn animals had increased glucose levels and other characteristics of MetSyn conditions, such as elevated levels of triglycerides and cholesterol ^9,37,38^. Further, we observed increased subcutaneous and visceral body fat with decreased muscle mass (Table 1 and Figure 1). Cross sectional analysis of the TA and soleus muscle from MetSyn animals showed left skewed in fiber type size distribution graph, indicating muscle atrophy (Figures 1C and D). The average cross-sectional area was significantly smaller in both MetSyn TA and soleus muscle (Figures 1A-D). The muscle atrophy in metabolic diseases is defined by the pathological term ‘sarcopenic obesity’ ^39–41^, and altered body composition signifies the important pathological aspects of our MetSyn model. In addition, the muscle loss in obesity or metabolic disease could result from chronic inflammation ^42–44^. Our previous data showed increased proinflammatory signaling in MetSyn mesentery bed with activation of M1 macrophages ^10^. In addition, dietary endotoxin, LPS, that also cause insulin resistance, altered inflammatory immune response in the lymphatic mesenteric bed ^45^. Hence, we speculate that muscle atrophy in fructose diet-induced MetSyn rats could be due to systemic inflammatory effects. Cardiac hypertrophic remodeling is one of the prevailing pathological features of metabolic disorders due to chronic systemic inflammation ^46–48^. Our data showed that hearts from fructose-induced MetSyn rats displayed increased heart weight and enlarged cardiomyocyte cross-sectional area (Table 1 and Figure 1).

SERCA is one of the key regulators of striated and smooth muscles’ contractions by regulating intracellular Ca^2+^ levels ^49,50^. While the role of SERCA in cardiac, skeletal, and smooth muscle have been largely investigated, very little is known about the role of SERCA in lymphatic muscle. One study showed that a SERCA inhibitor, cyclopiazonic acid (7μm) completely blocked lymph flow in bovine mesentery lymph vessels at all different transmural pressures ^25^. However, whether the inhibitory effects of SERCA was due to chronotropic or inotropic effects had not been tested. We employed thapsigargin that exhibits increased inhibitory effects on SERCA compared to other inhibitors ^51^. Our data showed that inhibition of SERCA significantly reduced lymphatic contractile frequency, amplitude and thus, diminished fractional pump flow and ejection fraction (Figure 2). These results imply that SERCA is an important mediator for lymphatic pumping by regulating phasic contractile activity, not by regulating lymphatic tonic contraction.

Diminished SERCA activity in metabolic diseases has been well established in various tissues, including cardiac, skeletal and smooth muscles, and in liver tissue. Consistent with our previous findings ^9,10^, lymphatic contractility indexes, frequency of contraction and lymphatic fractional pump flow, were significantly reduced in MetSyn rats when compared to control group (Figure 3C and E). Our data showing that the SERCA inhibitor, thapsigargin significantly reduces the contractility indexes of lymphatic vessels from control group, but not the vessels from MetSyn animals (Figures 3C, E and F), suggesting that SERCA activity is already diminished in MetSyn lymphatic vessels. Ejection fraction was significantly lowered in the presence of thapsigargin compared to each cohort groups (Figure 3D) and this might be due to the similar level of contractile amplitude between control and MetSyn.

Three distinct genes encoding SERCA 1, 2, and 3 produce more than 10 isoforms that are expressed in various muscles and non-muscle cells ^52^. SERCA1a and b are predominant isoforms in skeletal muscle and SERCA3s are expressed in various non-muscle tissues. SERCA2a is predominantly expressed in both skeletal muscle type 1 fiber and in adult cardiac muscle; while, SERCA2b is considered one of the major isoforms in smooth muscle cells ^52,53^. Here, we report that lymphatic muscle cells express both SERCA2a and SERCA2b isoforms (Figures 4A, 5D, F and G), further supporting our previous finding that lymphatic muscle presents a unique combination of muscle cell types that express both cardiac and smooth muscle contractile and regulatory proteins ^12,17–19^.

Previous studies in different metabolic disease models have shown a decreased SERCA activity in cardiac and vascular tissues, yet there were inconsistent results whether diminished SERCA activity resulted from decreased levels of SERCA expression or were independent of SERCA levels. Decreased SERCA2a protein expression was found in cardiac muscle coupled with impaired cardiac contractility in diabetic cardiomyopathy and in db/db mice ^28,29,54^. Additionally, SERCA2a protein levels decreased in vascular smooth muscle cells from Type 1 and Type 2 diabetes animal models ^27,55^. In contrast, studies have reported that there were conformational changes, altered SERCA regulatory molecules, or diminished SERCA activity without decreased SERCA expression in the cardiac tissues from db/db mice and in an insulin resistant rat model ^56,57^. We found that SERCA2a expression was diminished significantly in MetSyn lymphatics whereas SERCA2b expression was similar between control and MetSyn groups (Figures 4A-D). Thus, the decrease response to thapsigargin in MetSyn lymphatics that we observed could be due to the decrease levels of SERCA2a in lymphatic vessel. Decrease SERCA pump activity in lymphatic muscle would increase the cytosolic Ca^2+^ in LMCs. Our cell culture data support this notion; results showed that in the basal condition, intracellular Ca^2+^ levels were elevated in insulin resistant LMCs. Additionally, amplitude between peak and basal calcium levels was significantly decreased in insulin resistant LMCs, indicating decreased SERCA activity in LMCs treated with glucose and insulin. Increase in intracelluar Ca^2+^ will promote MLC_20_ phosphorylation via Ca^2+^/calmodulin-MLCK pathway ^22^. Increased MLC_20_ levels we observed in lymphatic vessels from MetSyn rats (Figures 4 E and F) corroborate with our previous finding that insulin resistant condition induced MLC_20_ phosphorylation in LMCs^21^. Thus, we propose that increased intracellular Ca^2+^ due to decreased SERCA2 expression and activity in LMCs of MetSyn animals lead to an increase in MLC_20_ phosphorylation that causes a reduction in the diameter of lymphatic vessels and consequently, poor lymphatic function in MetSyn conditions that we previously reported ^9,10^. Further studies are warranted to determine what isoforms of SERCA are present in lymphatic endothelial cells and if they play any roles in the modulation of flow-mediated Ca^2+^ release in MetSyn animals (Jafarnejad *et al.*, 2015).

Increasing SERCA expression or activity has been shown to provide protective effects on cardiac muscle and vascular smooth muscle cells in several metabolic diseases ^26,29,54,55^. A small molecule, CDN1163, activates SERCA by allosteric mechanism and improves Ca^2+^ homeostasis both *in vivo* and *in vitro* ^34–36^. Our data show that SERCA activation significantly improves the lymphatic contractile frequency in MetSyn lymphatic vessels similar to control lymphatic vessels (Figure 6 F). Though the fractional pump flow was not significantly improved by SERCA activator in MetSyn lymphatic vessels, fractional pump flow was not significantly lower than control lymphatic vessels, suggesting partial improvement of lymphatic contractile activity of MetSyn animals by SERCA activator.

In summary, the current study demonstrates that SERCA2 activation plays an important role in modulating lymphatic pump function and lymph flow by regulating chronotropic and inotropic effects. The rat mesenteric lymphatic muscle expresses both SERCA2a and SERCA2b isoforms. MetSyn conditions decreased the levels of SERCA2a expression and impaired Ca^2+^ regulation in LMCs that are coupled with increased MLC_20_ phosphorylation. The impaired lymphatic pumping activity in MetSyn is due to diminished SERCA activity and activating SERCA partially improves lymph flow in MetSyn. Therefore, it is possible that SERCA2 agonist could be used as a therapeutic strategy in enhancing lymphatic function in MetSyn or other metabolic diseases.

## METERIALS AND METHODS

### Materials

Phospho MLC_20_ (Ser19), total MLC_20_, β-actin antibodies were purchased from Cell Signaling Technology (Danvers, MA, USA). SERCA2a and SERCA2b antibodies were purchased from Badrilla Ltd (Leads, UK). Fura-2am, Fura-2am calibration kit, Alexa Fluor secondary antibodies, DMEM/F12, DMEM, fetal bovine serum (FBS), triple antibiotics (penicillin, streptomycin, and Amphotericin B), Prolong Gold antifade mounting medium with DAPI, WGA dye, and Alexa Flour 488 antibody labeling kit were purchased from Thermo Fischer Scientific (Waltham, MA, USA). SEC61A antibody was purchased from Abcam (Cambridge, MA, USA). DMSO, thapsigargin, CDN1163, bovine serum albumin, and α-SMA antibody were purchased Sigma Aldrich (St Louis, MO, USA). All other chemicals and reagents were from Sigma Aldrich (St. Louis, MO, USA), otherwise we indicated specifically.

### Animal Handling

Fifty-two male Sprague-Dawley rat weighing 150-180g were ordered from Charles River for induction of the MetSyn. Twenty rats were given a high fructose diet (60% fructose, ID89247 Harlan Teklad, Envigo, Indianapolis, IN, USA), for 7 to 10 weeks to induce the MetSyn, while the remaining rats were given standard rodent chow. Water and each respective feed were available ad libitum. Rats from the control group were utilized for testing the effects of different dosage of thapsigargin, DMSO, and CDN1163, a SERCA activator (n=12). Rats from the control and MetSyn group were used for isobaric functional analysis of mesenteric lymphatic vessel contractility (n=14/group). The remaining 6 rats were used for IHC and RNA analysis. All animals were housed in a facility with 12-h light-dark diurnal cycle, accredited by the Association for the Assessment and Accreditation of Laboratory Animal Care and maintained in accordance with the policies defined by the Public Health Service Policy for the Humane Care and Use of Laboratory Animals. All protocols had been approved by the University Laboratory Animal Compliance Committee at Texas A&M University prior to the commencement of the study.

### Determination of metabolic parameters

High fructose diet generates a rat model of MetSyn as characterized by high insulin, glucose, triglycerides, and glucose levels along with impaired insulin sensitivity ^37,58–60^. We also confirmed the development of MetSyn conditions by measuring plasma insulin, triglyceride, inflammatory cytokines in mesentery and cecum histopathological changes ^9,10^. To confirm the pathology of high fructose diet-induced MetSyn we measured blood glucose using a glucometer and test strips before surgery. Heart was isolated, weighted, and then the left ventricle was dissected for measuring cardiac hypertrophy. Quadriceps, tibial anterior (TA), and soleus muscles were isolated and weighed for assessing sarcopenic obesity, which is systemic pathology due to metabolic disorders. Inguinal subcutaneous adipose tissue (SWAT) and epididymal white adipose tissue (EWAT) were dissected to determine the percent of subcutaneous and visceral fat.

### Lymphatic vessel isolation and functional analysis

Rat mesenteric collecting lymphatic vessel isolation and cannulation were performed to test isobaric lymphatic pumping activity as described in our previous studies ^9,10,18^. Rats were anesthetized with Innovar-Vet (0.3 ml/Kg), which is a combination of a droperidol-fentanyl solution (droperidol 20mg/mL, fentanyl 0.4mg/ml), and diazepam (2.5mg/kg) intramuscularly. A midline excision was made and a loop of ileum was carefully exteriorized. Lymphatic vessels were carefully dissociated from the surrounding adipose tissue to prevent excess bleeding. Vessels were maintained in DMEM/F12 media at 38°C. Mesenteric lymphatic vessels were cannulated onto matched pipettes in a CH-1 chamber^®^ (Living Systems Instrumentation, St, Albans, VT, USA) and attached to separate pressure reservoirs. Vessels were incubated in DMEM/F12 or DMSO vehicle contained DMEM/F12 until it reached temperature and equilibrated at pressure 3 cmH_2_O for 30 minutes. After the equilibration, the vessel was recorded for 5 min at 1 cm, 3 cm, and 5 cm H_2_O of pressure. Vessels were incubated in the presence of DMSO (10-30mM), thapsigargin (1.0-2.0μm), or CDN1163 (1-10μm). At the end of each experiment, the bath solution was replaced with calcium-free albumin physiological saline solution and maximal diameter at each pressure recorded. Lymphatic diameters were traced for each 5 min video capture with vessel wall-tracking software developed and provided by Dr. Michael J. Davis at the University of Missouri-Columbia ^61^. Briefly, outer lymphatic vessel diameters were tracked 30 times per second, providing a tracing of diameter changes throughout the periods of systole and diastole. The following analogies to the cardiac pump parameters were derived: contraction amplitude, ejection fraction, contraction frequency, fractional pump flow, stroke volume, tonic index, lymph flow, and systolic/diastolic diameters as previously described ^17,62^.

### LMC culture and treatments

Primary rat mesenteric LMCs were obtained from mesenteric tissue explants of male Sprague-Dawley rats, as we have described in previous publications ^12,21^. Cells from passages 3-6 were used in all experiments. LMCs were cultured in DMEM, containing 10% FBS, and 1% triple antibiotics and maintained at 37 °C in 10% CO_2_ incubator. LMCs were plated in 24-well culture plates and then grown to ~70-80% confluence. The cells were serum starved for 24 h and treated with 5mM glucose (control) or high glucose (HG; 25mM), with or without insulin (100nM) for 48h as we reported in a previous study ^21^. To test the effects of free intracellular Ca^2+^, different concentrations of pCa solutions (between pCa7.5 to 3.5) were prepared based on Ca free DMEM using a software, MaxChelator as described ^63,64^. After serum starvation, LMCs were treated with Ca^2+^ free DMEM and different concentrations of free calcium (pCa 7.5-3.5) for 30 minutes. Proteins were isolated as described in our previous studies ^21,45^.

### Intracellular Ca^2+^ measurement in LMCs

Intracellular Ca^2+^ levels were measured using fura-2AM in phenol-free DMEM media as described ^65–67^. LMCs were plated in glass bottom chamber and treated when LMCs reached 60% confluence. Cells were loaded with Fura-2AM (2μM) in the dark for 30 mins at 37 °C. Cells were then washed with phenol-free DMEM and incubated for another 30 min for de-esterification. Pairs of fluorescent images were taken by exposure to 340- and 380nm double excitation with interference filters at selected wave length (Lamda DG-5, Sutter Instruments, Novato, CA, USA) and 510nm emission wavelengths using an epifluorescence microscopy system (Nikon Eclipse Ti, Nikon, Melville, NY USA). The individual traces of fura-2AM dye from multiple LMC region of interest, ROI were simultaneously measured from a field of view (~20 ROIs/plate). The fura-2 fluorescence ratio was collected for each cell throughout the experiments (NIS-Elements software, Nikon, Melville, NY, USA). Background fluorescence was determined before the start of the experiment. Background fluorescence was subtracted and fluorescence ratios of images at 340 and 380nm were determined. Calcium levels were measured using fura- 2AM calcium imaging calibration kit (Thermo Fisher) where it had zero to 10.0mM CaEGTA (1.35μM free calcium buffer) prior and post experiments. The interrelationship of the free Ca^2+^ concentration and the ratio was calculated according to manufacturer’s guideline ^68^. Basal Ca^2+^ levels were measured in phenol free-DMEM. Voltage-gated calcium channel were activated by depolarization with high (80mM) K^+^ solution. SERCA activity was assessed with thapsigargin (5μm).

### Immunofluorescence analyses

To determine the relative levels of SERCA2a, SERCA2b, and MLC_20_ phosphorylation, rat mesenteric lymphatic vessels were prepared for immunofluorescence as indicated in previous studies ^18,45^. Multiple adjacent lymphatic vessels were isolated and fixed in 2% paraformaldehyde for 60 mins at room temperature. After fixation, vessels were washed in PBS and permeabilized for 5 min in −20°C methanol. Vessels were then blocked in 5% goat serum (with 1% BSA) for 1 hour, subsequently cut in half with one part being used for the experimental treatment and the other part serving as the negative control. Vessels were incubated with SERCA2a (A010-23, Badrilla, Leads, UK), SERCA2b (A010-24, Badrilla, Leads, UK), α-Smooth muscle actin (A2547, Sigma-Aldrich, St. Louis, MO, US), or phospho MLC_20_ (3645, Cell signaling, Beverly MA, US) antibody in blocking solution overnight. Vessels were then washed and incubated with host matched secondary antibody (1:200) conjugated with Alexa Fluor®488 or Alexa Fluor ®594 after washed three times. Vessels were then washed three times and mounted using ProLong gold antifade mounting media. Vessels were scanned in 0.5-μm z-axis steps using PLAPLSM 40x objective (NA=0.9), Olympus IX-71 inverted microscope with Fluoview 300 confocal scanning. Average projections of series sections were reconstructed and quantified using Image J software ^69^. Negative control vessel segments were subjected to the same procedures as the experimental treatments except that negative controls were incubated with corresponding host IgG instead of the primary antibody. Negative controls were scanned at the same microscope settings as the experimental treatments to allow for valid comparison of the relative fluorescent intensities.

### WGA staining

Membrane specific dye, wheat germ agglutinin (WGA), was used to test MetSyn induced myopathies as described elsewhere ^70^. Frozen OCT block of TA, soleus, and left ventricles were cut 10 micron thickness using cooled cryostat (Leica CM1850, Leica Biosystems Inc., Buffalo Grove, IL, USA) at −20 °C. Sections were fixed in cooled acetone (−20 °C) for an hour. The slides were then washed three times in PBS for 5 minutes and incubated with WGA for 10 minutes at room temperature. Sections were washed and mounted using ProLong gold antifade mounting media. Images were taken with Olympus BX41 fluorescence microscope using UPlanApo x10 (NA=0.4) or UplanApo X20 objective (NA=0.7) (Olympus America, Melville, NY, USA) at an excitation peak of 545nm with an emission spectral peak of 610nM.

### ER and SERCA2a staining in LMCs

Immunofluorescence experiments in LMCs were performed as described earlier, with some modification ^21^, to determine SERCA2a expression and localization on ER. Briefly, the treated LMCs were fixed with 2% paraformaldehyde and permeabilized with cooled methanol for 4 mins at 4°C. Cells were then blocked with 1% BSA with 5% goat serum for 1 hour at room temperature. Cells incubated SERCA2a antibody (1:100) in blocking buffer for two hours at room temperature. After three washes, cells were incubated with Alexa Fluor 594 (goat anti-rabbit, 1:200) for 1 hour at room temperature in the dark. Since SEC61, specific to ER protein antibody is also from the rabbit, we used commercially pre-labeled SEC61 antibody (ab205794, Abcam, Cambridge, MA, USA), generated with an antibody labeling kit (A20181, Thermo Fisher, Waltham, MA, USA) and applied after SERCA2a antibody. Cell were counterstained for the nucleus and mounted using Prolong Gold antifade mounting medium with DAPI. Images were taken with Olympus BX41 fluorescence microscope using UPlanFLN X40 objective (NA=1.3) (Olympus America, Melville, NY, USA) and SERCA2a fluorescence intensity was measured with Image J.

### Protein isolation and Western blot analysis

Protein expression was quantified by Western blot analysis ^21,45^. In brief, LMC protein lysates were prepared from LMCs and separated onto a 4-20% gradient SDS-polyacrylamide gel and transferred, and Western blot analysis was performed. SERCA2a, SERCA2b, MLC_20_ phosphorylation, MLC_20_, and β-actin (1:1000) were used. Membranes were incubated with the appropriate horseradish peroxidase-conjugated secondary antibodies. Protein detection was conducted with an enhanced chemiluminescence system and visualized via image processor Fuji LAS-4000 Mini (GE Healthcare Bioscience, Pittsburg, PA, USA). β-actin expression was used as the loading control. Densitometric analyses was carried out using Image J (National Institute of Health, NIH, Bethesda, MD, USA). For quantification, experiments were repeated three times for each sample, and the resulting means ± SE were calculated (n=3/group).

### Statistical analysis

Animal characteristics (e.g., bodyweight, glucose level, % fat, muscle weight, CSA, etc.,) were analyzed by independent Student’s t-tests after performing Leven’s test for equal variance. Different dose effects of thapsigargin and CDN1163 in lymphatic contractility parameters were analyzed using One-way ANOVA with Dunnett’s post hoc test to find the treatment dose differences compared to control. Control and MetSyn lymphatic contractile parameters in the presence or absence of thapsigargin or CDN1163 were analyzed using Two-way ANOVA with Bonferroni’s post hoc test to examine the interaction and main effects between two factors: diet and treatment (i.e., thapsigargin,CDN1163). When we found a main effect or interaction, we performed follow-up ANOVA using Fischer’s LSD post hoc test to find between (i.e., diet) or within group (i.e., thapsigargin/CDN1163). Levene’s statistics also performed to test the homogeneity of variances. LMC data including Ca^2+^ were analyzed using Two-way ANOVA with Bonferroni’s post hoc test (glucose and insulin) as described earlier. Once we find any interaction or main effects, we performed follow-up ANOVA using Fisher’s LSD test. The homogeneity of variance was performed using Levene’s statistics. Statistical analyses was performed using SPSS software (IBM Corp, Armonk, New York, USA). All values are expressed as mean ± SE. The significance level was set at p<0.05. All graphs were generated with Prism 5 (GraphPad Software, La Jolla, CA, USA).

## Acknowledgements

The authors thank the Texas A&M Health Science Center Integrated Microscopy and Imaging Laboratory and Histology core facility at the department of Medical Physiology. We thank David Zawieja, PhD and his lab members, Wei Wang, MD and Olga Gasheva, MD, and Anatoliy Gashev, MD, PhD. for assisting with lymphatic functional studies. We also thank Dr. Cristine Heaps, PhD, and Jeffery Bray for technical assistant for calcium measurements. The authors thank Drs. Cynthia J. Meininger and Brett M. Mitchell for advice and proofreading. This work was supported by U.S. National Institutes of Health, National Institute of Diabetes and Digestive and Kidney Diseases Grant RO1 DK99221 (to M.M.) and Department of Medical Physiology, Lymphatic Graduate Student Fellowship (to Y.L).

## Author contributions

Y.L., S.C., and M. M. designed the study; Y.L. and S.C. performed the experiments. Y. L., S.C., and M.M. analyzed data and wrote the paper. All authors have approved submission of the final version of the manuscript. All authors agree to be accountable for all aspects of the work. All authors qualify for authorship and all those who qualify for authorship are listed.

## Data availability

The datasets analyzed during the current study are available from the corresponding author on reasonable request.

